# Social memory in female mice is rapidly modulated by 17β-estradiol through ERK and Akt modulation of synapse formation

**DOI:** 10.1101/2022.10.25.512385

**Authors:** Paul A. S. Sheppard, Deepthi Chandramohan, Alanna Lumsden, Daniella Vellone, Matthew C. S. Denley, Deepak P. Srivastava, Elena Choleris

## Abstract

**Background:** Social memory is essential to the functioning of a social animal within a group. Estrogens can affect social memory too quickly for classical genomic mechanisms. Previously, 17β-estradiol (E2) rapidly facilitated short-term social memory and increased nascent synapse formation, these synapses being potentiated following neuronal activity. However, what mechanisms underlie and co-ordinate the rapid facilitation of social memory and synaptogenesis are unclear. Here, the necessity of extracellular signal-regulated kinase (ERK) and phosphoinositide 3-kinase (PI3K) signaling for rapid facilitation of short-term social memory and synaptogenesis was tested.

**Methods:** Mice performed a short-term social memory task or were used as task-naïve controls. ERK and PI3K pathway inhibitors were infused intra-dorsal hippocampally 5 minutes before E2 infusion. Forty minutes following intrahippocampal E2 or vehicle administration, tissues were collected for quantification of glutamatergic synapse number in the CA1.

**Results:** Dorsal hippocampal E2 rapid facilitation of short-term social memory depended upon ERK and PI3K pathways. E2 increased glutamatergic synapse number (GluA1/bassoon colocalization) in task-performing mice but decreased synapse number in task-naïve mice. Critically, ERK signaling was required for synapse formation/elimination in task-performing and task-naïve mice, whereas PI3K inhibition blocked synapse formation only in task-performing mice.

**Conclusions:** Whilst ERK and PI3K are both required for E2 facilitation of short-term social memory and synapse formation, only ERK is required for synapse elimination. This demonstrates previously unknown, bidirectional, rapid actions of E2 on brain and behaviour and underscores the importance of estrogen signaling in the brain to social behaviour.

## Introduction

To behave appropriately within their social groups, social species require specialized cognitive abilities. Perhaps paramount of these is social recognition – the ability to recognize a conspecific or distinguish between conspecifics(1,2). Without social recognition, an animal can display maladaptive social behaviours due to inability to distinguish, for example, groupmates from intruders(3). Social behaviours, including social recognition, are processed through a “social brain network” including the medial extended amygdala, lateral septum, and certain hypothalamic nuclei(4,5). Importantly, while not thought of as a classically “social” brain region, the dorsal hippocampus is essential to the formation of social memories via inputs from social brain regions, and its disruption results in impairments in social memory(6–9). While much is understood with regards to hippocampal contributions to non-social memory (particularly spatial memory)(10), the integration of social information that occurs within the hippocampus requires further investigation.

Being reproductively active has been linked with social memory. Female mice show improved performance on a social recognition task during proestrus, the high estrogen and progesterone phase of the estrous cycle when females are sexually receptive(11). Ovariectomy results in impairments in social memory that can be rescued through administration of estrogens(12,13). Estrogens have long been known to elicit their effects by binding to intracellular estrogen receptors (ERs) that then dimerize, translocate to the nucleus, and directly affect gene transcription and protein expression(14). In addition to these delayed “classical” effects, estrogens affect molecular(15,16), cellular(17–19), systems(20,21), and behavioural(22–26) processes very rapidly (minutes) through intracellular mechanisms including activation of cell signaling cascades, such as the extracellular signal-regulated kinase (ERK) and phosphoinositide 3-kinase (PI3K) cascades.

Previous research discovered and richly characterized rapid effects of estrogens on short-term(27–29) and long-term memory in various tasks, such as spontaneous object recognition and object placement spatial memory tasks(25), as well as social memory(7,27–30). Given pre-acquisition in a 40-minute social recognition task, systemic(27), intra-dorsal hippocampal(7,28), and intra-medial amygdalar(30) administration of 17β-estradiol (E2), the most abundant and bioactive estrogen in adult mammals, facilitates short-term social memory in ovariectomized (OVX) female mice. The same doses of E2 increase hippocampal dendritic spine density in task-naïve mice(7,27) and in *ex vivo* hippocampal slices(28) within the same timeframe, consistent with other findings(19,31,32). However, the necessity of these spine increases to behaviour has been questioned(33). While an increase in dendritic spine number suggests an increase in synapse number, evidence indicates the relationship is more complex. For instance, frequency of AMPA miniature post-synaptic currents in *ex vivo* CA1 is reduced following the same treatment of E2 which increases CA1 dendritic spine density(28). Importantly, neuronal activity is needed for the formation of stable synapses from “silent” or nascent synapses(34,35). Ergo, synapse formation by estrogens, and not dendritic spine formation alone, may be a more salient and parsimonious mechanism through which estrogens affect behaviour. Synapse formation via priming by E2 and subsequent activation during learning may underlie the previously observed effects of E2 on short-term social memory.

Previous investigations into the mechanisms underlying the rapid effects of estrogens on memory have explored consolidation and long-term memory of non-social memories(25). However, short-term memory is necessary for the dynamic modulation of ongoing behaviours and needs to be processed in short periods of time(36). While some short-term memories will be consolidated into long-term memories, most information is not needed long-term, and the brain is selective in what memories are stored long-term. For instance, social information obtained during a party that adaptively modulates social responses for the duration of the party may become irrelevant in the days and weeks following. In view of their differing behavioral significance, it is perhaps unsurprising that the molecular underpinnings of short-term and long-term memory do not fully overlap (e.g. (37,38) but see (39,40) for discussion). The rapid and transient creation by estrogens of neuronal substrate for memory encoding provides a highly dynamic mechanism for short-term memory processing, especially in information rich, dynamic environments such as social interactions.

Estrogen-induced increases in synapse density observed both *in vivo*(19) and *in vitro*(17,34) suggest that pre-synaptic input to estrogen-treated neurons may be required for estrogen-mediated memory enhancements, as depicted in the concept of “two-step wiring plasticity”. This model posits that estrogens first transiently increase dendritic spine density, creating the neuronal substrate for learning to occur(34). Following activation (for instance, through memory task performance), these spines mature into persisting synapses(35). These estrogen-driven neurophysiological changes depend upon activity of cell signaling cascades, including ERK(16,17,34) and PI3K(17) pathways, which are also required for enhancement of object and spatial long-term memory consolidation by estrogens in female mice(32,41–45). While ERK is known to be necessary for the rapid increase in CA1 dendritic spine density following E2(32), the contributions of this kinase and the PI3K pathway to synapse formation *in vivo* are unknown. Additionally, two-step wiring plasticity suggests that the effects of estrogens on memory may depend upon activity (i.e. experience), leading to the hypothesis that learning events are drivers or modifiers of estrogen-mediated synaptic plasticity. However, whereas rapid effects of estrogens in dorsal hippocampus spine and synapse formation have been frequently reported(33), the underlying mechanisms remain unknown, especially in relation to their established beneficial roles in short-term social memory(7,28).

The present investigation explores the cellular mechanisms involved in E2 rapid facilitation of short-term social memory. We investigate whether the facilitating role of E2 in the dorsal hippocampus on short-term social memory requires activation of the ERK or PI3K cell signaling cascades. Additionally, we investigate interplay between the rapid effects of E2 and short-term social memory task performance on CA1 glutamatergic synapse formation.

## Methods

Detailed methods can be found in Supplementary Information.

### Subjects

Young adult (2-month-old), experimentally naïve female CD1 mice (*Mus musculus*) were used (Charles River, Kingston, NY, USA). Following surgeries, experimental mice were single-housed for 10-15 days prior to experiments. Stimulus mice were single-housed for 7 days post-surgery then pair-housed for a minimum of 7 days prior to participating in behavioural testing. All behavioural tests were run during the dark cycle (between 09:00h and 19:00h) in the experimental animals’ home cages under red light. All procedures were approved by the University of Guelph Animal Care and Use Committee and followed the guidelines of the Canadian Council on Animal Care.

### Surgeries

All mice were ovariectomized (OVX) to minimize gonadal hormone levels and fluctuations. Within the same surgical session, experimental mice were further implanted with bilateral guide cannulae directed at the dorsal hippocampus. Stimulus mice were OVX to ensure that their hormone status would not affect investigative behaviours by experimental mice.

### Rapid Social Recognition Paradigms

Two rapid social-recognition paradigms of short-term social memory were used **[Figure 1A]**. The first, “easy” paradigm was designed such that vehicle-treated OVX mice show social recognition by preferentially investigating a novel over a previously encountered social stimulus at test(29). OVX mice receiving treatment that impairs social recognition will show no preference between the 2 stimuli. In this “easy” paradigm, experimental mice were exposed to 2 novel OVX stimulus mice for three 4-minute sample phases **[Figure 1B]**. Sample phases were separated by 3-minute rest periods in which no stimuli were present in the cage. After the final sample phase and 3-minute memory retention period, 2 stimuli were reintroduced into the cage for a 4-minute test phase: one novel stimulus and one previously encountered stimulus from the sample phases **[Figure 1B]**.

**Figure 1:**
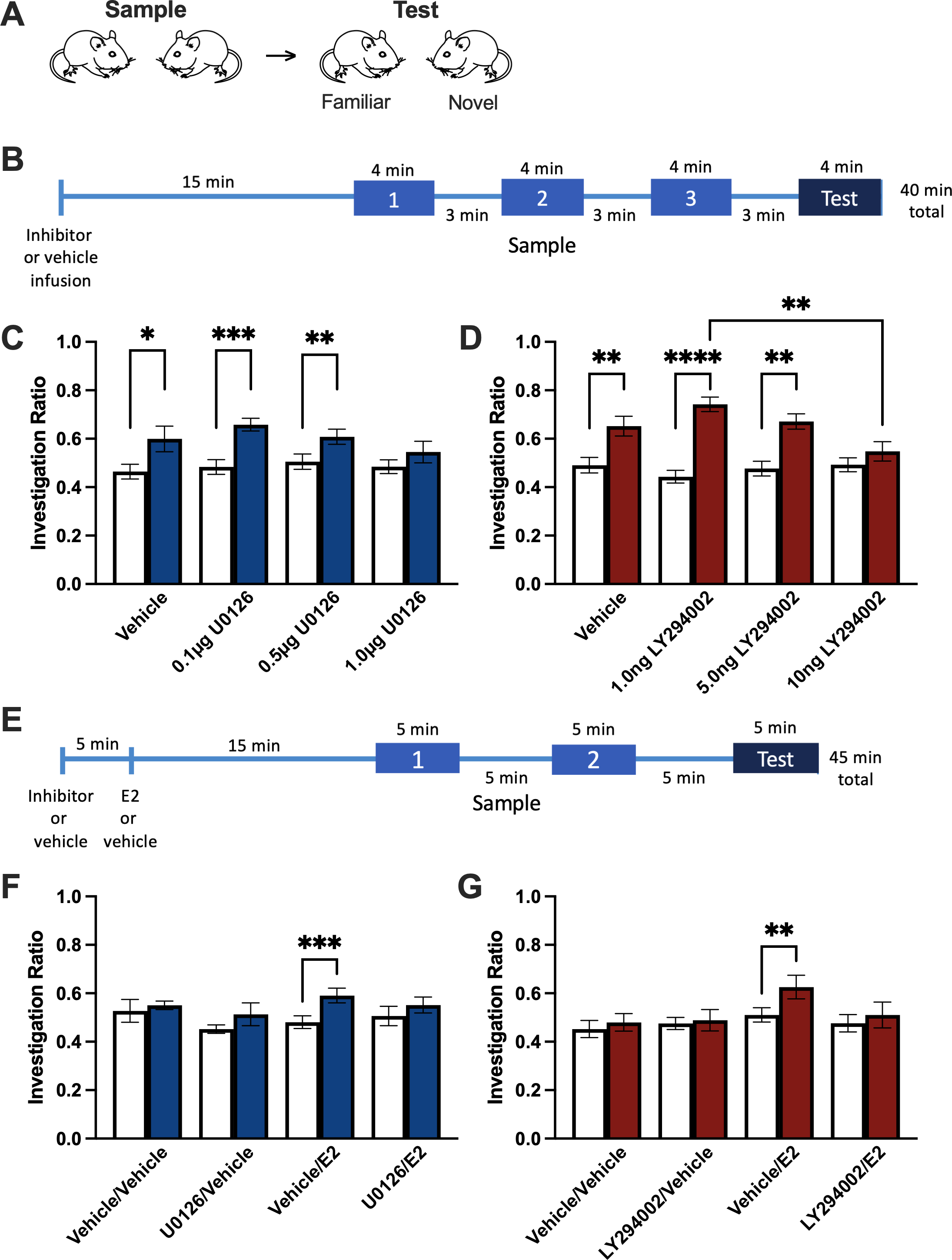
Rapid effects of 17β-estradiol on short-term social recognition are blocked by ERK and PI3K pathway inhibition. A) Schematic of overall short-term social recognition testing. B) Timeline of treatment and “easy” short-term social recognition testing. C) Intra-dorsal hippocampal U0126 (ERK pathway inhibitor, 1.0μg/side) blocks short-term social recognition memory. D) Intra-dorsal hippocampal LY294002 (PI3K pathway inhibitor, 10ng/side) blocks short-term social recognition memory. E) Timeline of treatment and “difficult” short-term social recognition testing. F) Intra-dorsal hippocampal U0126 blocks the facilitating effects of E2 on short-term social recognition memory. G) Intra-dorsal hippocampal LY294002 blocks the facilitating effects of E2 on short-term social recognition memory. * p<0.05, ** p<0.01, *** p<0.001, **** p<0.0001. Data presented as mean ± SEM.

To show facilitating effects of treatment, a “difficult” rapid social recognition paradigm, similar to the “easy” version except that there are two 5-minute sample phases and one 5-minute test each separated by 5-minute rests **[Figure 1E]**, was used(29). The decreased number of exposures to the stimulus mice makes this task more difficult than the “easy” paradigm and vehicle-treated OVX CD1 mice do not exhibit short-term social memory, whereas those who receive treatments that facilitate memory for the social stimulus (e.g. E2) do(7,27–30,46,47).

### Treatment administration

#### Effects of ERK or PI3K pathway inhibition on social recognition

OVX mice were bilaterally microinfused (0.5μL/side at 0.2μL/minute) with 0.1, 0.5, or 1.0 μg/side of MEK inhibitor 1,4-diamino-2,3-dicyano-1,4-bis[2-aminophenylthio]butadiene (U0126; Promega, Madison, WI); 0.5, 1.0, 5.0, or 10ng/side of PI3K inhibitor 2-(4-morpholinyl)-8-phenyl-4H-1-benzopyran-4-one (LY294002; Santa Cruz Biotechnology, Dallas, TX); or vehicle (50% dimethyl sulfoxide [DMSO] in 0.9% NaCl solution) and then tested on the “easy” social recognition paradigm 15 minutes following the beginning of the infusion **[Figure 1B]**. In all experiments, infusers were left in place for an additional minute following each infusion to ensure the full dose was administered and to prevent back-flow.

#### Effects of ERK or PI3K pathway inhibition on estradiol-facilitated social recognition

OVX mice were bilaterally microinfused first with 0.5μg/side U0126, 5.0ng/side LY294002, or vehicle (0.25μL/side at 0.2μL/minute) 5 minutes before 6.81pg/side 17β-estradiol (E2; Sigma-Aldrich, Oakville, ON, Canada) or vehicle (0.25μL/side at 0.2μL/minute) and then tested on the “difficult” social recognition paradigm 15 minutes following the beginning of the infusion **[Figure 1E]**. This dose of E2 previously facilitated social recognition in OVX female mice in the “difficult” paradigm(7,28). These doses of U0126 and LY294002 did not impair social recognition in OVX female mice in the “easy” paradigm **[Figure 1C and D]**.

#### Effects of estradiol and cell signaling inhibition in task-naïve OVX mice

OVX mice were bilaterally microinfused with the same treatments as in estradiol-facilitated social recognition experiments above (vehicle, 0.5μg/side U0126, or 5.0ng/side LY294002 followed by vehicle or 6.81pg/side E2) and left undisturbed for 40 minutes before tissue collection to determine the effects of treatment on synapse number in the dorsal CA1 of task-naïve mice.

### Immunohistochemistry, confocal imaging, and analysis

Coronally sectioned hippocampus from task-performing and task-naïve groups (4-6 mice/treatment) were used for immunohistochemistry, Briefly, sections were permeabilized in PBS supplemented with 0.05% Triton-X100; blocked in 10% Normal Goat Serum, 1.5% BSA, 0.3% Triton-X100 in PBS. for 3-4 hours> Primary antibodies: Rabbit-α-GluA1 (Sigma-Aldrich AB1504, 1:300) and Mouse-α-bassoon (Abcam ab82958, 1:200) were incubated overnight in blocking solution. Sections were then incubated in secondary antibody diluted in blocking solution (1:1000 Goat-α-rabbit AlexaFluor488; 1:1000 Goat-α-mouse AlexaFluor568) for 2 hours, before coverslipping with mounting medium containing DAPI (Prolong Gold; Thermofisher). For antigen retrieval, sections were incubated for 10-15 minutes in 10mM sodium citrate (pH 6.2) at RT, followed by a 15-minute incubation with 10mM sodium citrate (pH 6.2) at 78°C before permeabilization.

Confocal images of the CA1 region using an Inverted Spinning Disk confocal microscope (Nikon, Japan) and 60x oil immersion lens objective (NA 1.4). Exposure time was kept constant for the entire dataset. Images were acquired as a stack spanning 6-10μm, at an interval of 0.3μm. Three 3 non-continuous slices were imaged and analysed from each animal, with data averaged to a single datum for each animal. Synaptic puncta were analysed in ImageJ (https://imagej.net/Welcome), using a previously published pipeline(48). Analysis of synaptic puncta was performed in the strata oriens and radiatum – corresponding to the basal and apical dendritic regions respectively of CA1 pyramidal neurons **[Figure 2A]** and limited to 50×100μm Region of interest (ROI) 20μm either side of the stratum pyramidale to limit synaptic analysis to secondary and higher dendritic branching **[Figure 2A]**. following parameters described in (48). In all sections, GluA1 or bassoon expression was first determined. Synaptic puncta were determined by assessing the number of GluA1 puncta that overlapped with bassoon puncta.

**Figure 2:**
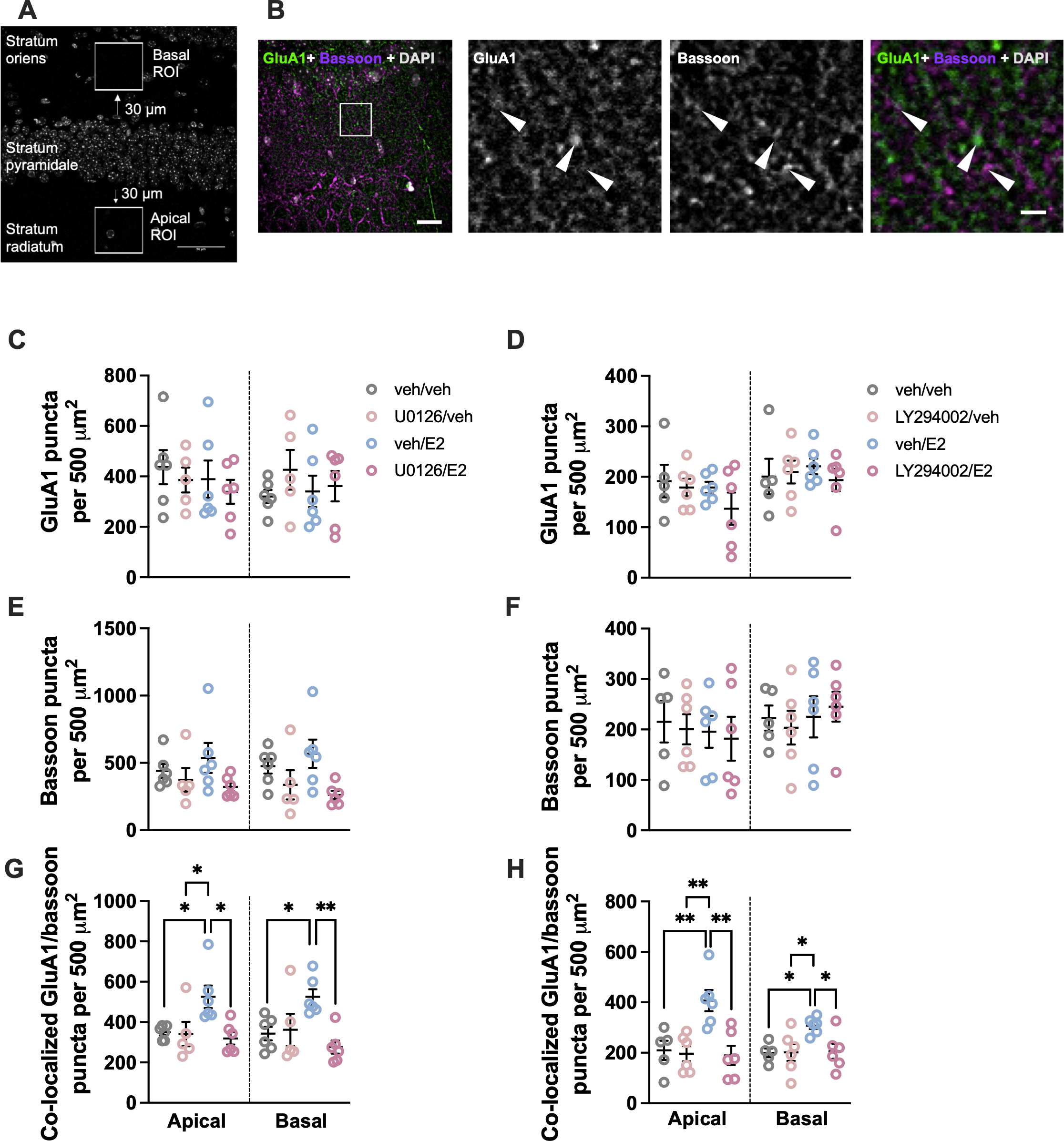
17β-estradiol rapidly increases glutamatergic synapse number in CA1 pyramidal neurons in short-term social recognition task-performing mice. A) Schematic depicting apical and basal ROIs in dorsal CA1 slices. B) Representative images of GluA1 puncta, bassoon puncta, and GluA1/bassoon colocalization. C-D) GluA1 puncta density is unaffected by intra-dorsal hippocampal E2 or inhibitor (U0126 [C] and LY294002 [D]) treatment. E-F) Bassoon puncta density is unaffected by intra-dorsal hippocampal E2 or inhibitor (U0126 [E] and LY294002 [F]) treatment. G) E2 increases glutamatergic synapse number in task-performing mice. ERK pathway inhibition blocks this effect. H) E2 increases glutamatergic synapse number in task-performing mice. LY294002 pathway inhibition blocks this effect. * p<0.05, ** p<0.01. Data presented as mean ± SEM.

### Behaviour data analysis

Videos were collected from all sample and test phases and analyzed (The Observer, Noldus Information Technology, Wageningen, The Netherlands) for both social (sniffing stimuli, digging/burying near stimuli, etc.) and non-social (horizontal movement, vertical non-investigative behaviour, grooming, etc.) behaviours(7,27–30,46,47). Active sniffing within 1-2mm of a stimulus mouse-containing cylinder was considered social investigative behaviour. An investigation ratio was calculated; IR = N/(N+F), in which N is the time spent investigating the novel (in sample phases, N is the stimulus that will be replaced) and F is the time spent investigating the familiar stimulus.

Investigation ratios for sample phases were averaged for analysis.

### Statistical analysis

The arcsin transformed investigation ratios were analyzed with a mixed-design repeated measures ANOVA with treatment as the main factor and paradigm session (average sample and test) as the repeated-measure dependent variable. To reduce type I errors, specific mean comparisons were planned *a priori* to assess differences between IR_Sam_ and IR_Test_ within each treatment group. One-way ANOVAs were used to assess treatment effects on IR_Test_, followed by Tukey *post hoc* tests. The durations of the other behaviours (Supplementary Table 1) were analyzed using a mixed-design repeated measures ANOVA with treatment as the main factor and paradigm session (each of the sample phases and test) as the repeated-measures dependent variable, followed by Tukey *post hoc* tests. When normality failed, Kruskal-Wallis ANOVAs were performed followed by Dunn’s *post hoc* tests. One-way ANOVAs with Tukey *post hoc* tests were used to determine effects of treatment and task performance on GluA1 and bassoon puncta and GluA1/bassoon colocalization. SPSS and GraphPad Prism (v9.3.0) were used for all statistical analyses. Cohen’s d and eta-squared effect sizes are provided where appropriate. Two-tailed statistical significance was set at p<0.05.

## Results

### ERK and PI3K inhibition in the dorsal hippocampus blocked short-term social memory in a dose-dependent manner

To evaluate the necessity of the cell signaling cascades to E2-facilitated short-term social memory, we first determined whether and at what doses intra-dorsal hippocampal ERK or PI3K inhibition blocked short-term social memory in an “easy” task where control animals can discriminate familiar from novel conspecifics. As indicated by a significant difference between sample and test investigation ratios, vehicle-treated control mice (t=2.56, df=9, Cohen’s d=0.991, p=0.031), as well as 0.1μg/side (t=4.72, df=14, Cohen’s d=1.66, p=0.0004) and 0.5μg/side (t=3.49, df=12, Cohen’s d=0.937, p=0.0051) U0126-treated mice had significant short-term social memory, whereas 1.0μg/side U0126 impaired short-term social memory (p=0.0932; main effect of session: F_(1,42)_=37.41, η^2^=0.201, p<0.0001)[**Figure 1C]**.

Similarly vehicle-(t=4.64, df=10, Cohen’s d=1.40, p=0.0012), 1.0ng/side (t=6.72, df=10, Cohen’s d=3.21, p<0.0001), and 5.0ng/side (t=4.57, df=10, Cohen’s d=1.89, p=0.001) LY294002-treated mice had significant short-term social memory, whereas 10ng/side LY294002-treated mice did not (p=0.118; main effect of session: F(1,39)=82.68, η^2^=0.375, p<0.0001; session by treatment interaction: F(3,39)=6.79, η^2^=0.0922, p=0.0009)**[Figure 1D]**. Additionally, there was a main effect of LY294002 treatment on IR_Test_ (F_(3,39)_=5.21, η^2^=0.286, p=0.004), with 10ng/side significantly lower than 1.0ng/side LY294002 (p=0.002)**[Figure 1D]**. These effects are consistent with and comparable to effects seen on long-term object recognition memory(41,42). Total social investigation time was unaffected between groups **[Figures S1-2]**. Therefore, short-term social memory is impaired by either ERK or PI3K pathway inhibition, implicating these pathways as necessary for short-term social memory.

### Dorsal hippocampal ERK or PI3K inhibition block E2-facilitated short-term social memory

Having determined the doses at which ERK and PI3K pathway inhibitors block short-term social memory, we determined whether intra-dorsal hippocampal E2 requires ERK and/or PI3K signaling to facilitate short-term social memory. Therefore, the highest doses of U0126 (0.5μg/side) and LY294002 (5ng/side) that did *not* block social memory in the “easy” social recognition task were used in conjunction with E2 (6.81pg/side), a dose known to facilitated short-term social memory in the ‘difficult’ task(7,28). E2-treated mice exhibited significant short-term social memory (t=4.20, df=15, Cohen’s d=1.01, p=0.0009 **[Figure 1F]**; t=3.31, df=11, Cohen’s d=0.858, p=0.0079 **[Figure 1G]**), replicating previous findings(7,28), whereas vehicle and inhibitor-only controls did not (ps>0.261, main effect of session [F_(1,46)_=8.43, η^2^=0.0592, p=0.0056]**[Figure 1F]**; ps>0.512, main effect of session [F_(1,43)_=5.85, η^2^=0.0294, p=0.020]**[Figure 1G])**. Critically, the facilitating effects of E2 were blocked by ERK (p=0.258) or PI3K inhibitor infusion (p=0.42)**[Figures 1F-G]**. E2-treated mice in the ERK-pathway experiment showed greater social investigation than vehicle controls (p=0.0363; F_(3,46)_=3.05, η^2^=0.111, p=0.0379)**[Figure S2]**. However, we do not see a similar effect in the PI3K-pathway experiment **[Figure S4]** nor in previous investigations(7,28). Furthermore, neither the E2-nor vehicle-treated groups differed from either U0126-treated group (ps>0.102), suggesting that this increase in social investigation by E2 does not explain treatment effects on short-term social recognition memory. Overall, these data show that signaling through both the ERK and PI3K pathways is required for the rapid facilitation of short-term social memory by E2.

### E2 increased glutamatergic synapses in task-performing mice in an ERK- and PI3K-dependent manner

As E2 required both the ERK and PI3K pathways to facilitate social memory, and knowing that E2 rapidly inhibits glutamatergic transmission in CA1 dorsal hippocampal pyramidal neurons of task-naïve mice(28) we investigated the effect of dorsal hippocampal E2 with or without pre-treatment with ERK or PI3K inhibitors on glutamatergic synapse formation following social recognition training. Treatment did not affect GluA1 or bassoon expression in either apical or basal sub-fields in task-performing mice (no main effects of treatment, ps>0.238 and ps>0.0525, respectively **[Figures 2C-F, 3A-D]**. However, E2-treated mice had a higher density of synaptic puncta (colocalization of GluA1 and bassoon puncta) than vehicle treated mice (apical – Cohen’s d=1.76, p=0.0365, main effect of treatment [F_(3,19)_=5.10, η^2^= 0.446, p=0.0093]; basal – Cohen’s d=2.08, p=0.0476, main effect of treatment [F_(3,19)_=5.42, η^2^=0.461, p=0.0076]**[Figure 2G]**; apical – Cohen’s d=2.10, p=0.0086, main effect of treatment [F_(3,19)_=8.13, η^2^=0.562, p=0.0011]; basal – Cohen’s d=3.02, p=0.0412, main effect of treatment [F(3,19)=4.36, η^2^=0.408, p=0.0170]**[Figure 2H])**. The effect of E2 on synaptic puncta in trained animals was blocked by pre-treatment with either U0126 (ps<0.0123) or LY294002 (ps<0.0441)**[Figures 2G-H]**. Conversely, treatment with these inhibitors alone had no effect on synaptic puncta (U0126: ps>0.993; LY294002: ps>0.995)**[Figure 2G-H]**. Together, these data show that E2 administration in task-performing mice increases glutamatergic synapse number and that this effect requires ERK and PI3K signaling.

**Figure 3:**
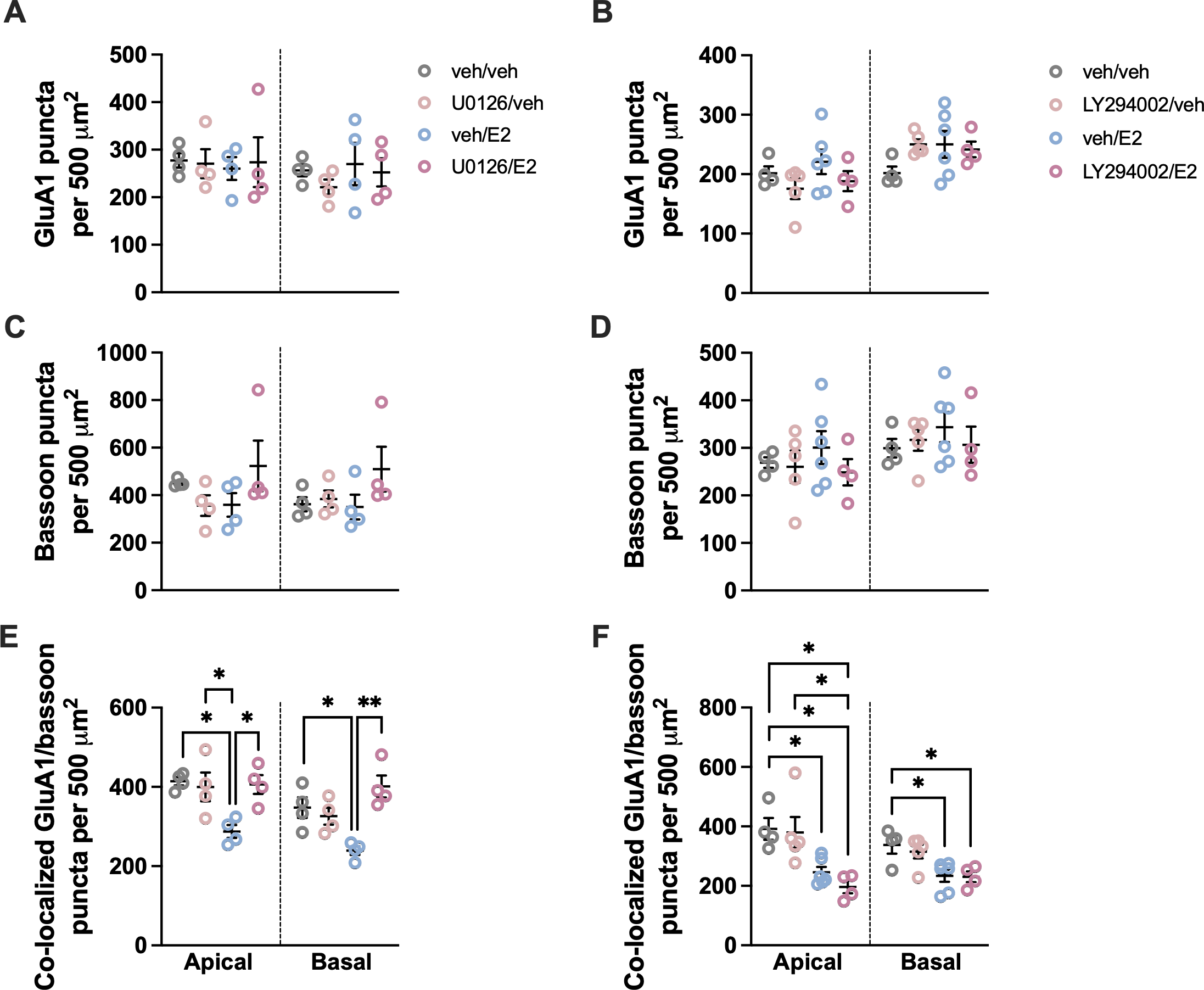
17β-estradiol rapidly decreases glutamatergic synapse number in CA1 pyramidal neurons in task-naïve mice. A-B) GluA1 puncta density is unaffected by intra-dorsal hippocampal E2 or inhibitor (U0126 [A] and LY294002 [B]) treatment. C-D) Bassoon puncta density is unaffected by intra-dorsal hippocampal E2 or inhibitor (U0126 [E] and LY294002 [F]) treatment. G) E2 increases glutamatergic synapse number in task-performing mice. ERK pathway inhibition blocks this effect. H) E2 increases glutamatergic synapse number in task-performing mice. LY294002 pathway inhibition blocks this effect. * p<0.05, ** p<0.01. Data presented as mean ± SEM.

### E2 decreased glutamatergic synapse number in task-naïve mice in an ERK-, but not PI3K-, dependent manner

To evaluate whether task-performance affects the effects of E2 on synapse formation, task-naïve mice receiving the same treatments as task-performing mice were sacrificed following the same delay but without performing the social recognition task. Analysis of GluA1 and bassoon expression in the CA1 of these mice revealed that E2 decreased synaptic puncta in both basal and apical dendrites of pyramidal neurons (apical – Cohen’s d=4.67, p=0.0121, main effect of treatment [F_(3,12)_=6.40, η^2^=0.616, p=0.0078]; basal – Cohen’s d=2.70, p=0.0233, main effect of treatment [F_(3,12)_=8.98, η^2^=0.692, p=0.0022]**[Figure 3E]**; apical – Cohen’s d= 2.42, p=0.0397, main effect of treatment [F_(3,15)_=7.40, η^2^=0.597, p=0.0029]; basal; Cohen’s d=1.92, p=0.0250, main effect of treatment [F(3,15)=5.70, η^2^=0.533, p=0.0083]**[Figure 3F]**), without an effect on the total expression of either synaptic protein (apical – ps>0.237, basal – ps>0.208)**[Figures 3A-D]**. This decrease was blocked by U0126 microinfusion prior to E2 (Cohen’s ds>2.92, ps<0.0183), but not by LY294002 (ps>0.746)**[Figures 3E-F]**. Synaptic puncta in mice that received LY294002 and E2 did not differ from the E2-only group, but was significantly reduced compared to vehicle controls (Cohen’s ds>2.23, ps<0.035)**[Figure 3F]**. This suggests that E2 decreases glutamatergic synapse number in task-naïve mice via ERK, but not PI3K, signaling.

## Discussion

Social cognition is essential for adaptive behaviours in social species for which rapid modification of behaviour is integral to responding to dynamic changes in the social environment. Critical to this is the ability to recognize and distinguish between conspecifics. It is now recognized that estrogens act as potent modulators of brain and behaviour, including social memory(7,27,28,30), within minutes(22,25). Recent evidence provides potential neuronal substrate for these effects: 17β-estradiol rapidly increases dendritic spine density(7,27,28), suggesting an increase in synapse formation, and modulates glutamatergic signaling(17,28,34), an effect that appears dependent upon synaptic plasticity. A causal link between E2 rapid effects on brain and behaviour has yet to be established. As such, we asked the questions 1) what intracellular processes are required for E2 facilitation of social memory?; 2) does E2 modulate synapse number in both social memory task-performing and task-naïve animals?; and 3) do the same intracellular processes required for E2’s modulation of social memory also co-ordinate E2-mediated synapse formation? As predicted by the two-step wiring plasticity model, the present results demonstrate rapid and dynamic modulation of short-term social memory and hippocampal glutamatergic synapse number by 17β-estradiol is dependent upon ERK and PI3K signaling and task-performance.

ERK and PI3K pathways in the dorsal hippocampus were both found to be required for short-term social memory per se and its facilitation by E2, similarly to long-term object and spatial memory(32,41–44). This novel finding suggests that similar intracellular mechanisms may be at play in short- and long-term memory, the parsimonious explanation for which being that the dorsal hippocampus, in both regards, is functioning as a crucial memory processing centre. While often regarded as a hub of spatial memory formation(10), accumulating evidence implicates the dorsal hippocampus in the processing of social information in interplay with upstream and downstream regions of the social brain(6–9,28), and here we show that the ERK and PI3K pathways are critical to social memory in this region. Estrogens modulate social cognition(2,49), including social memory. We have previously shown that systemic or intra-dorsal hippocampal administration of E2 rapidly facilitates short-term social memory and that these same treatments also increase dendritic spine density in CA1 pyramidal neurons of task-naïve mice(7,27,28). Here, we have elucidated key intracellular mechanisms underlying the rapid facilitation of short-term social memory by E2. ERK and PI3K were previously found to be rapidly activated by estrogens(16,41,42) and necessary for long-term memory enhancements by E2(32,41–43,45). Our results suggest that these pathways are also necessary for the rapid enhancement of short-term social memory, thus underlying the adaptive dynamic modulation of social interactions by estrogens at times of changing social milieu.

While the rapid effects of E2 on short-term social memory and synapse formation occur within the same timeframe, the causal link previously remained elusive. Here, E2 in the dorsal hippocampus rapidly facilitated short-term social memory and increased glutamatergic synapse number in the same task-performing animals. Furthermore, when facilitation of social memory by E2 was blocked by ERK or PI3K pathway inhibition, so too was synapse formation. This suggests that glutamatergic synapse formation drives the rapid facilitating effects of E2 on social memory and does so in an ERK- and PI3K-dependent manner. Remarkably, in task-naïve animals, E2 decreased synaptic puncta – that is, in the absence of task-performance, E2 decreases glutamatergic synapse number. Consistent with these findings, we have previously shown that E2 induced the formation of silent synapses(28) that can be potentiated by synaptic stimulation(34). There is a high degree of functional relevance to these novel findings. Memory formation requires formation of novel synapses, but unchecked synapse formation is likely maladaptive and would lead to interference between memory traces (50,51). Here, we have shown E2 to eliminate existing synapses in a paucity of stimulation (i.e. lack of task performance) and increase synapse number when a learning event occurs (i.e. performance of social recognition task), consistent with two-step wiring plasticity mechanism of estrogen action(35). This fine-tuning of estrogens’ rapid synaptic effects by task performance may sharpen the signal of relevant memory traces by increasing task relevant synapses and decreasing irrelevant synapses, thereby increasing the signal-to-noise-ratio. Furthermore, the data in this study demonstrate that the bi-directional effects of synapse density are driven by signalling through the ERK pathway – ERK inhibition blocked both E2-induced increases and decreases in GluA1/bassoon puncta in task-performing and task-naïve mice, respectively. Conversely, PI3K appears to have a more specific role – inhibition of this pathway blocked E2-mediated increases in synaptic puncta in task-performing mice but did not reverse E2’s ability to reduce synaptic density in task-naïve mice. Through these previously unknown bidirectional effects, E2 is capable of both providing plasticity and modifying its use.

Estrogens are potent neuromodulators in a variety of behaviours(23,26,52) and in learning and memory(22,25,53–55). Throughout the lifespan, estrogens allow for adaptive responses to the challenges brought on by a dynamic world(49), including dynamic social environments. Foundational to this are the rapid effects of E2 on short-term social memory and modulation of hippocampal glutamatergic synapse number described here. That there are bidirectional effects of E2 on synapse number dependent upon task-performance introduces a new dynamic neuromodulation of brain and behaviour by estrogens which warrants further future investigation.

## Supporting information

Supplementary Results

## Acknowledgements and Disclosures

Work performed at the University of Guelph was funded by the Natural Sciences and Engineering Research Council of Canada (NSERC; RGPIN-2018-04699 and RGPAS-522695-2018) and the Canada Foundation for Innovation (Grant No. 9585) to EC.

Work performed at King’s College London was supported by the Medical Research Council (MRC) Centre grant (MR/N026063/1). DPS also acknowledges funding from the Medical Research Council (MR/L021064/1) and Royal Society UK (Grant RG130856). He is also a recipient of an Independent Investigators award from the Brain and Behavior Foundation (formally National Alliance for Research on Schizophrenia and Depression (NARSAD); Grant No. 25957),

The authors thank Jenna Ashley and Marika Gummieny-Matsuo for help performing behavioural experiments and the Wohl Cellular Imaging Centre (WCIC) at the IoPPN, King’s College, London, for help with microscopy.

The authors have nothing further to disclose.

Data are available upon reasonable request to the corresponding author.

