## Supplementary Figures for "Social memory in female mice is rapidly modulated by 17β-estradiol through ERK and Akt modulation of synapse formation"

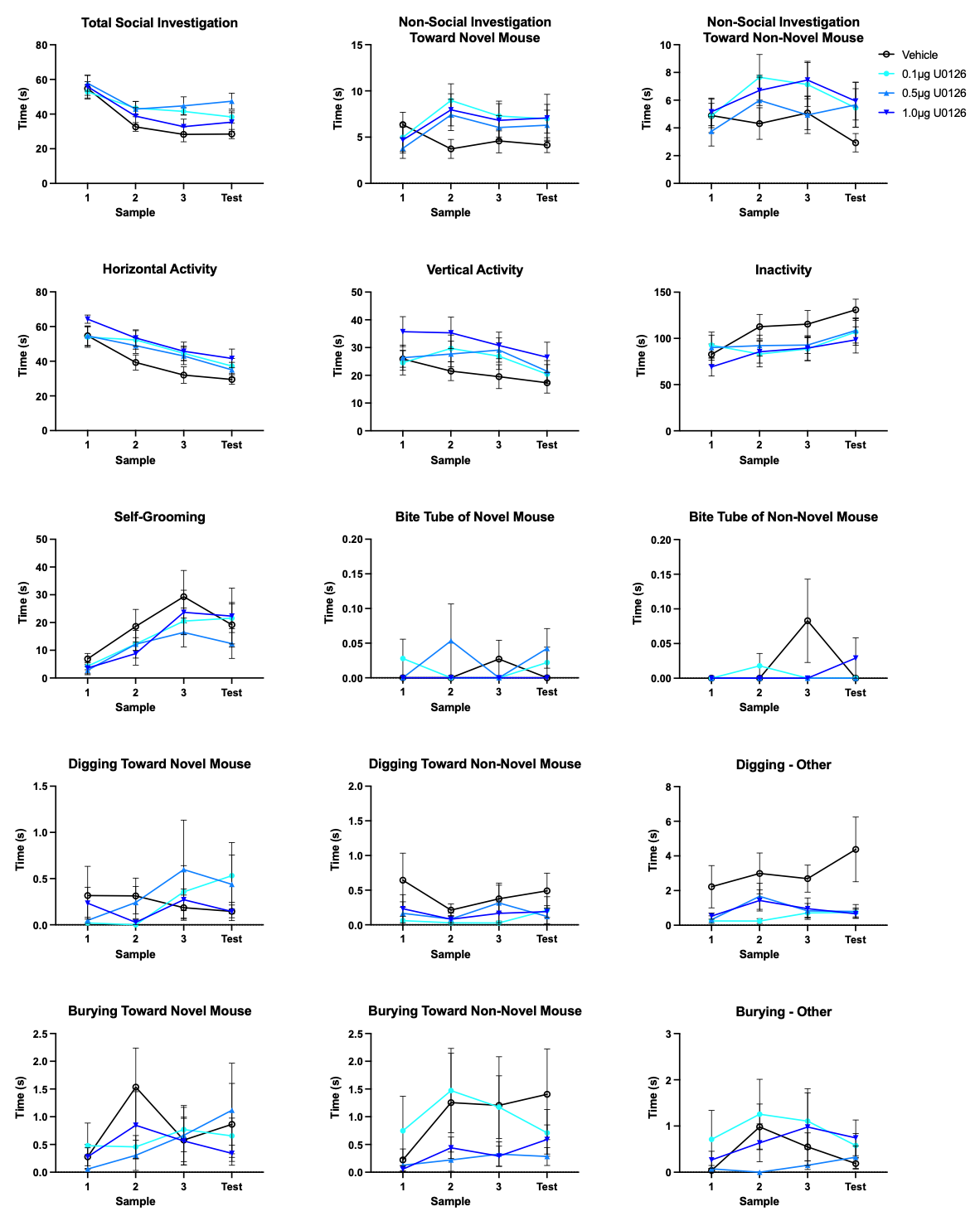

**Figure S1: Effects of ERK pathway inhibition on non-social behaviours in the “easy” short-term social memory task**

There was a significant treatment by session interaction in inactivity (F_(9,141)_=2.285, η^2^=0.0193, p=0.020), with no significant *post hoc* comparisons**.** Vehicle treated mice spent more time digging away from conspecifics than did 0.1µg/side U0126 mice (p=0.0252; F_(3,47)_=3.48, η^2^=0.126, p=0.023). Total social investigation time and other behaviours were unaffected. Data presented as mean ± SEM.

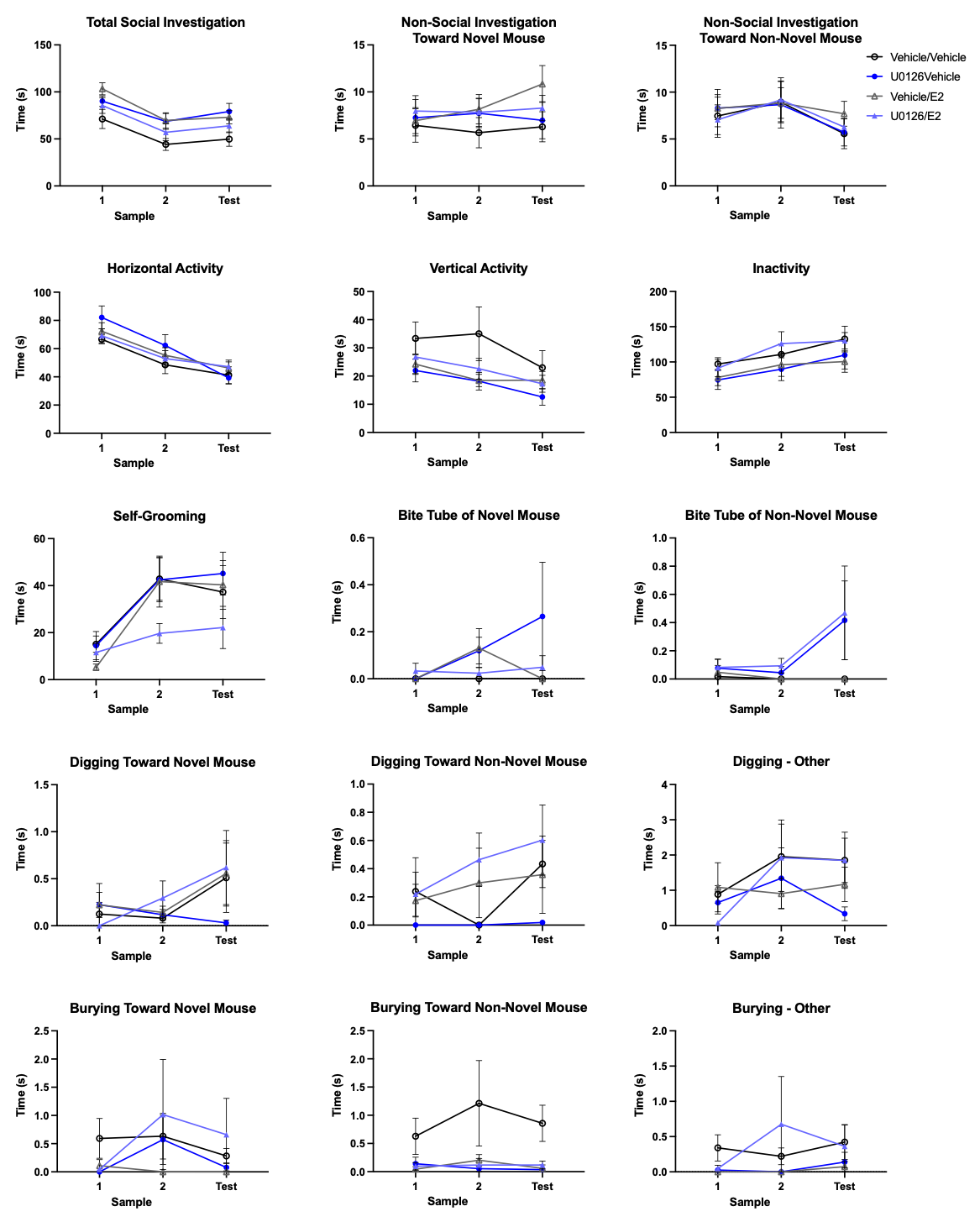

**Figure S2: Effects of ERK pathway inhibition and E2 on non-social behaviours in the “difficult” short-term social memory task**

Vehicle-treated mice spent more time burying toward the non-novel conspecific than all other groups (F_(3,46)_=5.44, η^2^=0.146, p=0.0028; ps<0.0095 for each pairwise comparison). Importantly, E2 treated mice showed greater social investigation than vehicle controls (p=0.0363; F_(3,46)_=3.05, η^2^=0.111, p=0.0379). However, neither the E2 nor the vehicle groups differed from either U0126 treated group (ps>0.102), suggesting that this increase in social investigation by E2 does not explain treatment effects on short-term social recognition memory. Other behaviours were unaffected. Data presented as mean ± SEM.

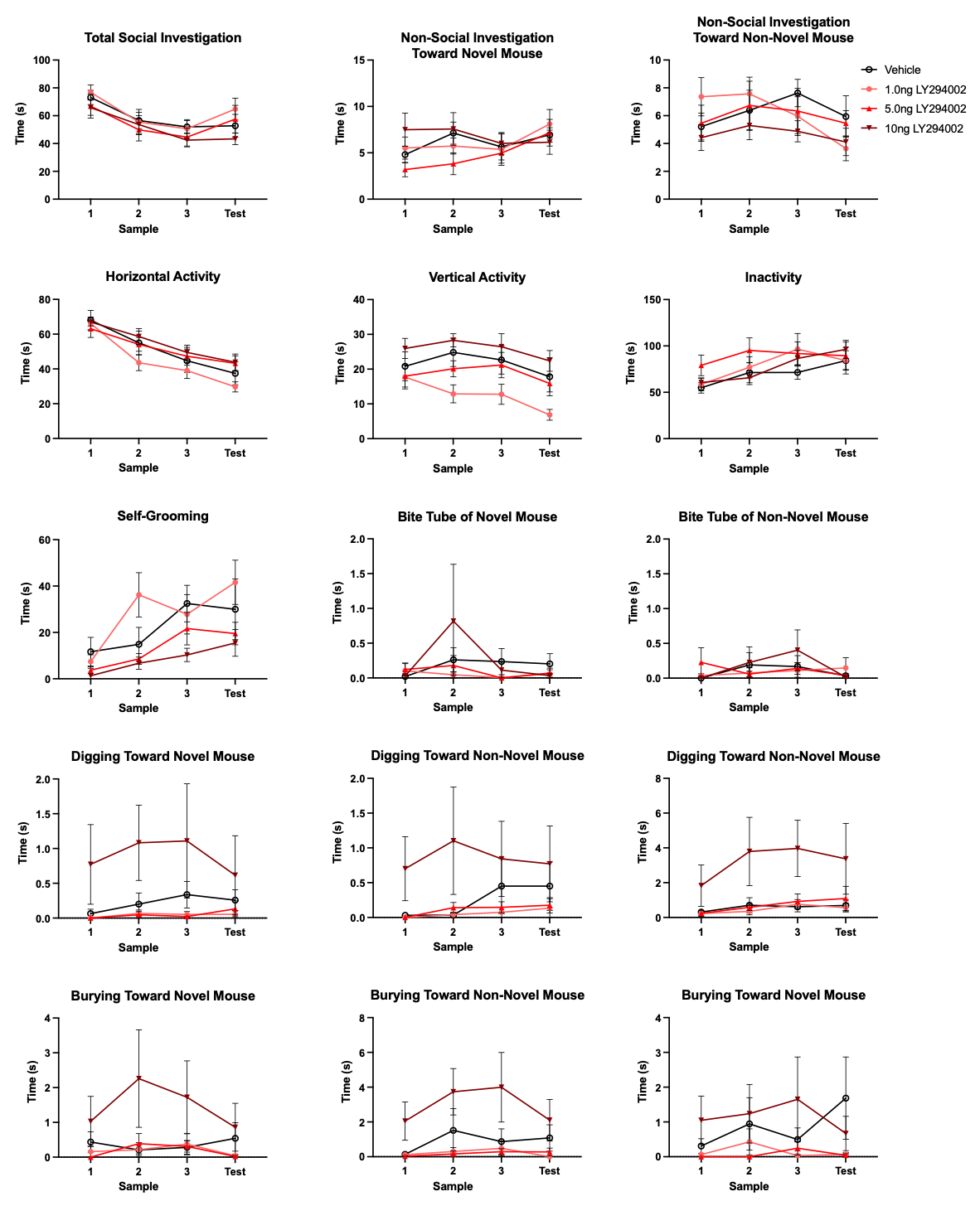
 **Figure S3: Effects of PI3K pathway inhibition on non-social behaviours in the “easy” short-term social memory task**

Mice given 10ng/side LY294002 exhibited more vertical activity than 1.0ng/side mice (p=0.0061; F_(3,39)_=4.30, η^2^=0.173, p=0.0103) and more burying towards the non-novel conspecific than 1.0ng/side (p=0.0457) and 5.0ng/side mice (p=0.0422; F_(3,39)_=3.36, η^2^=0.152, p=0.0283). Total social investigation time and other behaviours were unaffected. Data presented as mean ± SEM.

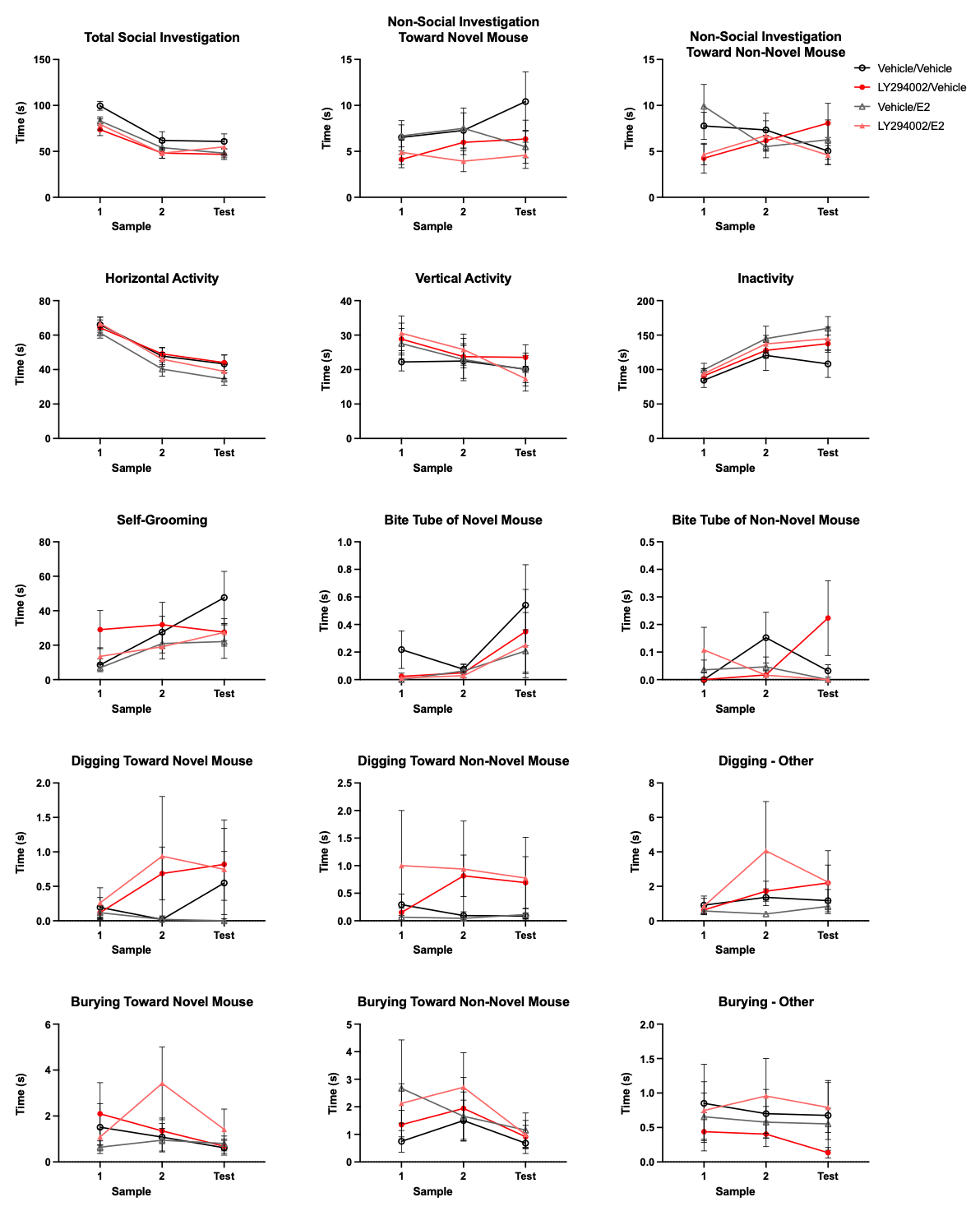

**Figure S4: Effects of PI3K pathway inhibition and E2 on non-social behaviours in the “difficult” short-term social memory task**

There were significant treatment by session interactions in non-social investigation (F_(6,36)_=2.86, η^2^=0.0641, p=0.0137, no significant *post hocs*) and biting (F_(6,36)_=2.90, η^2^=0.108, p=0.0127) the tube of the non-novel mouse, with *post hocs* of the latter revealing differences between LY294002 only treated mice and both E2-treated groups (E2 only: p=0.0332, LY294002 and E2: 0.0281). Total social investigation time and other behaviours were unaffected. Data presented as mean ± SEM.
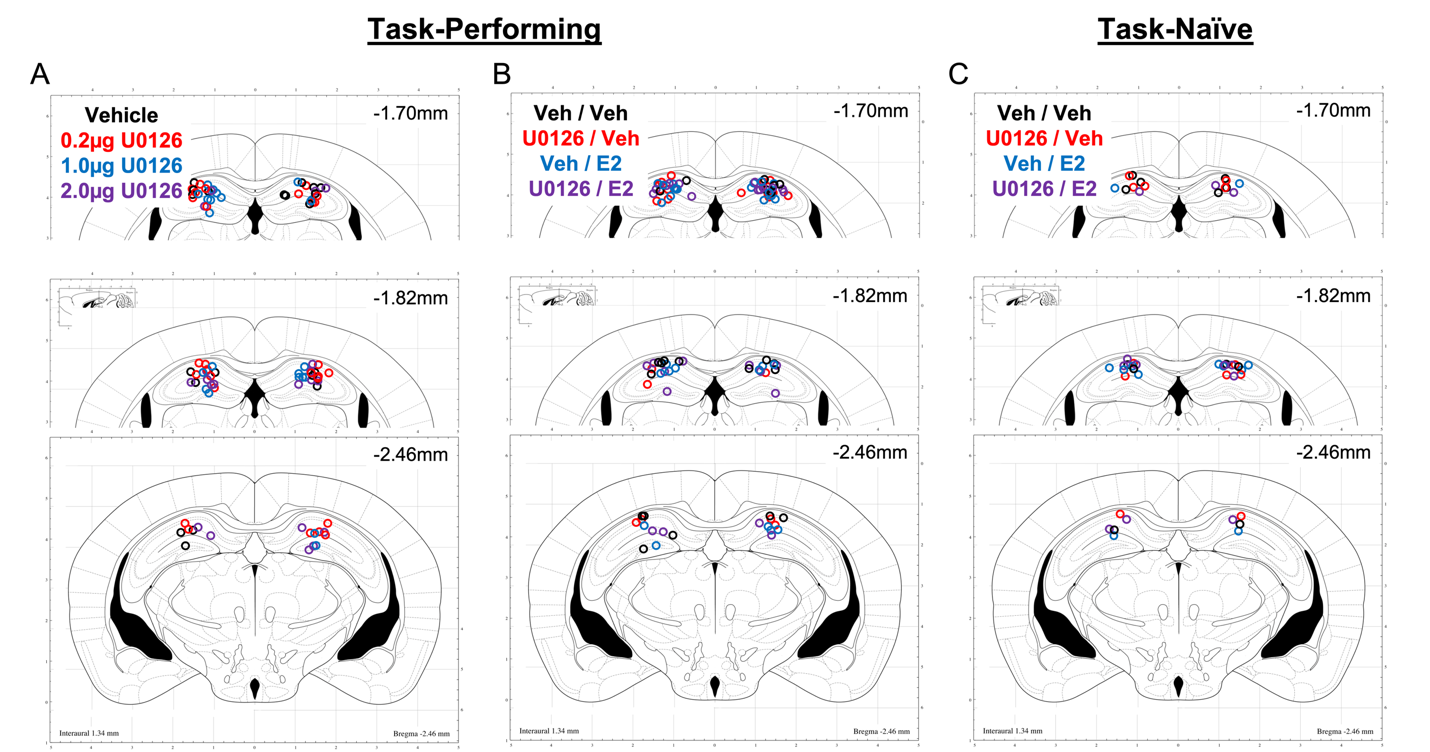
 **Figure S5: Cannula placements for ERK-pathway experiments**

Cannula placements for OVX mice implanted with intra-dorsal hippocampal bilateral guide cannulas. A) Placements for ERK pathway inhibition in “easy” social memory experiment. B) Placements for ERK pathway inhibition in “difficult”, E2-facilitated social memory experiment. C) Placements for task-naïve ERK pathway inhibition experiment. Only mice with the injectors in the dorsal hippocampus are shown. All other OVX mice were excluded from the behavioural and dendritic spine analyses.

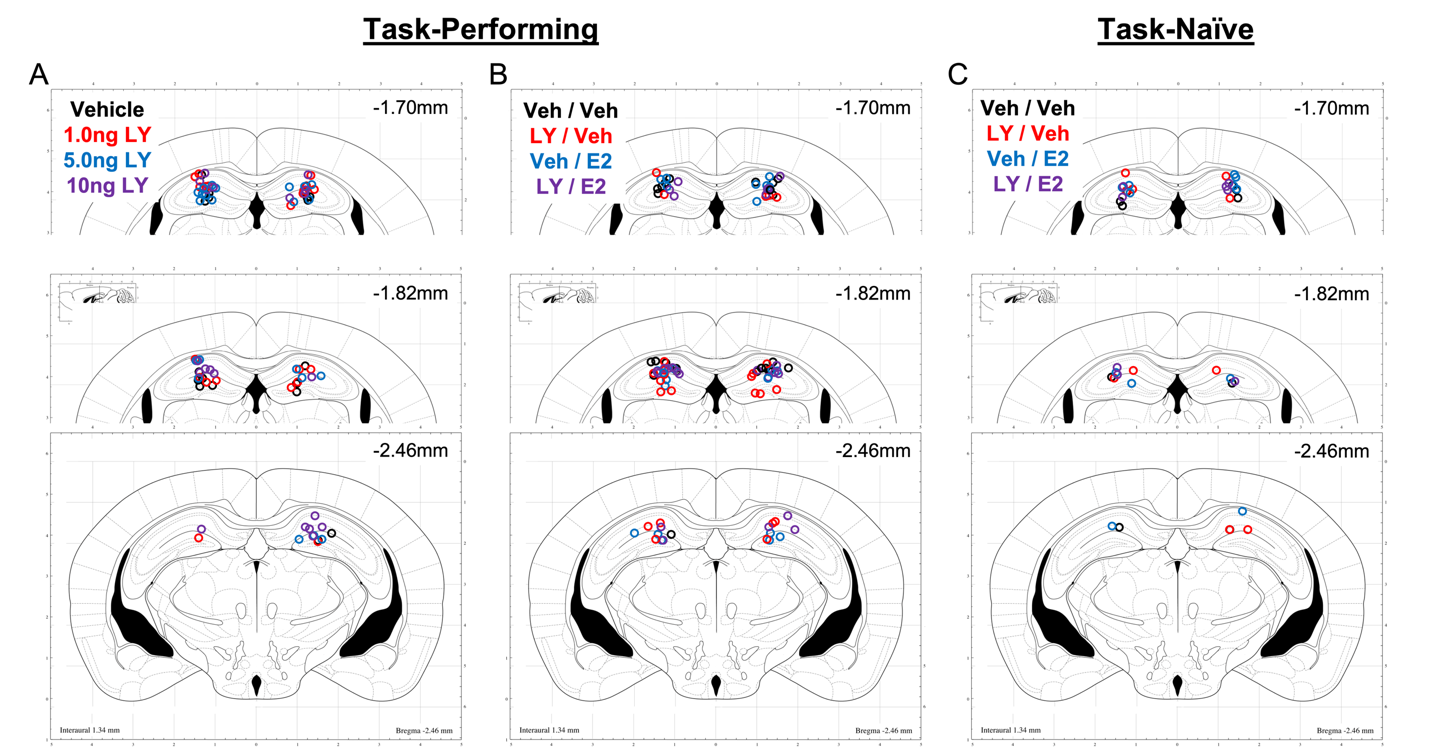
 **Figure S6: Cannula placements for PI3K-pathway experiments**

Cannula placements for OVX mice implanted with intra-dorsal hippocampal bilateral guide cannulas. A) Placements for PI3K pathway inhibition in “easy” social memory experiment; LY indicates LY294002. B) Placements for PI3K pathway inhibition in “difficult”, E2-facilitated social memory experiment. C) Placements for task-naïve PI3K pathway inhibition experiment. Only mice with the injectors in the dorsal hippocampus are shown. All other OVX mice were excluded from the behavioural and dendritic spine analyses.
