## Supplementary Methods for "Social memory in female mice is rapidly modulated by 17β-estradiol through ERK and Akt modulation of synapse formation"

**Subjects**

Two-month old (young adult), experimentally naïve female CD1 mice (*Mus musculus*) were used (Charles River, Kingston, NY, USA). Upon arrival, mice were initially triple-housed in clear polyethylene cages (26x16x12cm^3^) containing corncob bedding, environmental enrichment (paper nesting material, paper cups, and/or clear plastic houses), and *ad libitum* tap water and rodent chow (Teklad Global 14% Protein Rodent Maintenance Diet, Harlan Teklad, WI). Mice were given a minimum of 7 days to acclimate to the colony room before surgeries. Following surgeries, experimental mice were single-housed for 10-15 days before participating in any experiments. Stimulus mice were single-housed for 7 days post-surgery before being pair-housed for a minimum of 7 days prior to participating in behavioural testing. All animals were housed on a reversed light/dark cycle (12:12h, lights on at 20:00h) at 21±1°C with 40-50% humidity. All test mice were used in only one behavioural experiment. Stimulus mice were reused for 8-16 testing days. All behavioural tests were run during the dark cycle (between 09:00h and 19:00h) in the experimental animals’ home cages under red light. Animals were moved into the experimental rooms the evening prior to testing to acclimate. Vaginal smears were taken to confirm successful ovariectomies from vaginal cytology(1). All procedures were approved by the University of Guelph Institutional Animal Care and Use Committee and followed the guidelines of the Canadian Council on Animal Care.

**Surgeries**

All mice were ovariectomized (OVX) to minimize gonadal hormone levels and fluctuations. Within the same surgical session, experimental mice were further implanted with bilateral guide cannulae directed at the dorsal hippocampus. Stimulus mice were OVX to ensure that investigative behaviours by experimental mice were not affected by hormone status of the stimuli.

All mice were subcutaneously injected with the analgesic and anti-inflammatory drug carprofen (50mg/kg; Rimadyl, Pfizer Canada Inc, Kirkland, QC, Canada) at least 1 hour prior to being anesthetized with isoflurane (Benson Medical Industries, Markham, ON) and placed in a stereotaxic frame using atraumatic ear bars (David Kopf Instruments, CA). They then received 0.02mL of a local anesthetic mix of 0.17% bupivacaine (Hospira, Inc., Montreal, QC, Canada), 0.67% lidocaine (Alveda Pharmaceuticals, Toronto, ON, Canada), and saline solution (0.9% NaCl) subcutaneously into incision sites. Post-surgery, mice were administered 0.5mL of warm saline solution intraperitoneally to restore hydration.

***Ovariectomy surgeries*.** Each anesthetized mouse had the fur on its lower back shaved and the area cleaned with antiseptic solutions. A small dorsal incision of no more than 2cm was then made in the skin followed by two smaller (<1cm) bilateral lumbar incisions in the muscles overlying the ovaries. One at a time, the ovarian arteries and oviducts were clamped, the ovaries excised, and the remaining tissue inserted back into the incision. The skin incision was closed with 1-2 MikRon Autoclip 9mm wound clips (MikRon Precision Inc., Gardena, CA) and a few drops of local anesthetic were applied.

***Cannulation stereotaxic surgeries*.** Following ovariectomy, the dorsal head of each anesthetized experimental mouse was shaved and cleaned. The scalp was then excised and the membrane overlying the skull was sloughed off using 3% hydrogen peroxide (H_2_O_2_). Two holes were then drilled into the skull to accommodate the 26-gauge bilateral guide cannulae (Plastics One, HRS Scientific, Anjou, QC, Canada) just dorsal to the CA1 region of the dorsal hippocampus, at 1.7mm posterior to bregma, 1.5mm lateral to midline, and 1.3mm below the skull surface(2). Injectors (Plastics One, HRS Scientific, Anjou, QC, Canada) extended 1mm beyond the end of the guide cannulae, making the final injection depth 2.3mm below the skull surface. Three jeweller’s screws (Plastics One, HRS Scientific, Anjou, QC, Canada) were screwed into 3 additional drilled holes in the skull around the perimeter of the cannula pedestal. The cannulae were held in place with dental cement (Central Dental Ltd, Scarborough, ON, Canada) which completely covered the skull surface exposed in the procedure. Dummy cannulae (Plastics One, HRS Scientific, Anjou, QC, Canada) flush with guide cannulae were inserted.

**Rapid Social Recognition Paradigms**

Mice were gently restrained by hand to insert the infuser and then microinfused while moving freely using a microinfusion pump (PHD 2000, Harvard Apparatus, QC, Canada) and then returned to their home cage with environmental enrichment removed and a clear plexiglass cover replacing the metal feeding and drinking lid. To test the rapid effects of treatment on short-term memory, the social recognition paradigm was completed within 40 minutes of inhibitor **[Figure 1B]** or hormone **[Figure 1E]** administration. Experiments were recorded from above by Everio digital camcorders (HD Everio GZ-E300, JVC, Mississauga, ON, Canada).

Two rapid social-recognition paradigms of short-term social memory were used. The first, “easy” paradigm was designed such that vehicle treated OVX mice show social recognition by preferentially investigating a novel stimulus over a previously encountered stimulus at test(3). OVX mice receiving a treatment that impairs social recognition will show no preference between the 2 stimuli. In this “easy” paradigm, experimental mice were exposed to 2 novel OVX stimulus mice for three 4-minute sample phases (Figure 1B). Sample phases were separated by 3-minute rest periods in which no stimuli were present in the cage. After the final sample phase and 3-minute memory retention period, 2 stimuli were reintroduced into the cage for a 4-minute test phase: one novel stimulus and one previously encountered stimulus from the sample phases (Figure 1B).

To show facilitating effects of treatment, a “difficult” rapid social recognition paradigm was used (3). This social recognition paradigm is similar to the “easy” version except that there are two 5-minute sample phases and one 5-minute test each separated by 5-minute rests (Figure 1E). The decreased number of exposures to the stimulus mice makes this task more difficult than the “easy” paradigm and vehicle treated OVX CD1 mice do not show short-term social memory, whereas those who receive treatments that facilitate memory for the social stimulus (e.g. E2) do(3–9).

In both easy and difficult rapid social recognition paradigms stimulus mice were held in clear Plexiglas tubes (7cm diameter, 12cm high), to which they had been previously habituated, with perforations (36 holes, 4mm diameter) around the bottom third to allow olfactory and social cues to be transmitted to the test mouse, but to keep the stimulus mice in a constant location and from initiating interactions. These tubes were cleaned with Fisherbrand Sparkleen detergent and baking soda between uses and thoroughly dried to eliminate any olfactory traces of other mice.

**Treatment administration**

Mice were bilaterally infused with a cell signaling pathway inhibitor or vehicle at a rate of 0.2μL/minute to a volume of 0.5μL/side (2.5min) and then tested on the “easy” social recognition paradigm 15 minutes following the beginning of the infusion **[Figure 1B]** or bilaterally infused with a cell signaling pathway inhibitor or vehicle 5 minutes prior to E2 or vehicle **[Figure 1E]**, each infusion at a rate of 0.2μL/minute to a volume of 0.25μL/side (1.25min). The “difficult” social recognition paradigm began 15 minutes following the beginning of the second infusion. Task-naïve OVX mice were infused the same treatments as mice tested on the “difficult” paradigm and left undisturbed in their cages for 40 minutes before tissue collection, corresponding to the time when the mice performing the social task completed testing. Infusers were left in place for an additional minute following each infusion to ensure the full dose was administered and to prevent back-flow.

**Effects of ERK pathway inhibition on social recognition**

OVX mice were bilaterally microinfused with 0.1 µg/side, 0.5 µg/side, or 1.0 µg/side of MEK inhibitor 1,4-diamino-2,3-dicyano-1,4-bis[2-aminophenylthio]butadiene (U0126; Promega, Madison, WI) or vehicle (50% dimethyl sulfoxide [DMSO] in 0.9% NaCl solution). The highest dose (1.0 µg/side) was previously found to block long-term OR memory consolidation(10) and the middle dose (0.5 µg/side) blocked rapid increases in pERK following E2-treatment(11).

**Effects of ERK pathway inhibition on estradiol-facilitated social recognition**

OVX mice were bilaterally microinfused first with 0.5µg/side U0126 or vehicle 5 minutes before 6.81pg/side 17β-estradiol (E2; Sigma-Aldrich, Oakville, ON, Canada) or vehicle. This dose of E2 previously facilitated social recognition in OVX female mice in the “difficult” paradigm used here in which vehicle treated OVX mice do not show social recognition(5,6). This dose of U0126 did not impair social recognition in OVX female mice in the “easy” paradigm in which vehicle treated OVX mice show social recognition **(Figure 1C)**.

**Effects of PI3K pathway inhibition on social recognition**

OVX mice were bilaterally microinfused with 0.5ng/side, 1.0ng/side, 5.0ng/side, or 10ng/side of PI3K inhibitor 2-(4-morpholinyl)-8-phenyl-4H-1-benzopyran-4-one (LY294002; Santa Cruz Biotechnology, Dallas, TX), or vehicle (50% dimethyl sulfoxide [DMSO] in 0.9% NaCl saline solution). These doses were based on Fan et al. (12), where LY294002 at 5.0ng/side blocked long-term object recognition memory and at 0.5ng blocked rapid increases in phospho-PI3K following E2-treatment.

**Effects of PI3K pathway inhibition on estradiol-facilitated social recognition**

OVX mice were bilaterally microinfused first with 5.0ng/side LY294002 or vehicle 5 minutes before 6.81pg/side E2 or vehicle. This dose of LY294002 did not impair social recognition in OVX female mice in the “easy” paradigm in which vehicle treated animals show social recognition **(Figure 1D)**.

**Effects of estradiol and cell signaling inhibition in task-naïve OVX mice**

OVX mice were bilaterally microinfused with the same treatments as in estradiol-facilitated social recognition experiments above (vehicle, 0.5µg/side U0126, or 5.0ng/side LY294002 followed by vehicle or 6.81pg/side E2) and left undisturbed for 40 minutes before tissue collection to determine the effects of treatment on synapse number in the dorsal CA1 of task-naïve mice.

**Tissue Collection**

Immediately after social recognition testing, mice received 0.8mL of the anesthetic Avertin intraperitoneally (tribromoethanol; Sigma-Aldrich, Oakville, ON, Canada) and was perfused with 0.9% NaCl solution, followed by 4% paraformaldehyde (PFA) in phosphate buffered solution (PBS; pH 7.4) to fix the brain *in situ*. The brain was subsequently extracted and post-fixed in 4% PFA in PBS overnight before storage in PBS containing 0.02% sodium azide. Brains from each animal were serially sectioned (25 or 40 μm-thick coronal sections) on a [cryostat](https://www.sciencedirect.com/topics/neuroscience/cryostat) at −18 °C and stored in tissue cryoprotection solution (25 % glycerol, 30 % [ethylene glycol](https://www.sciencedirect.com/topics/neuroscience/ethylene-glycol), 45 % 1x PBS pH 7.4, 0.05 % sodium azide) at −20 °C until further processing. Coronal sections (40μm) were also generated and mounted on microscope slides to confirm location of cannulae (Supplemental Figures)(2). Mice were removed from analyses when infusion cannulae did not reach the dorsal hippocampus bilaterally (total n=8 across all experiments).

**Immunohistochemistry**

Coronal sections mounted on Superfrost Plus slides (ThermoFisher Scientific, UK), from each brain in task-performing and task-naïve groups (4-6 mice/treatment) were washed for 10 min in phosphate buffer (PB; 0.1 M) and 2 × 10 min in PBS. For antigen retrieval sections were incubated for 10−15 mins in 10 mM sodium citrate (pH 6.2) at RT, followed by a 15 min incubation in pre-heated 10 mM sodium citrate (pH 6.2) in a water-bath at 78 °C. Sections were then allowed to cool down to RT in the same solution while gently shaking for 30 min. Sections were then washed twice (2 × 5 min) in PBS supplemented with 0.05 % Triton-X100 and incubated for 3−4 h in blocking solution (10 % Normal Goat Serum, 1.5 % BSA, 0.3 % Triton-X100 in PBS), followed by overnight incubation at 4 °C with primary antibodies diluted in blocking solution for the GluA1-subunit of AMPA receptors and the pre-synaptic marker bassoon: 1:300 Rabbit-α-GluA1, Sigma-Aldrich AB1504; 1:200 Mouse-α-bassoon, Abcam ab82958). Sections were then washed in PBS (3 × 10 min) and incubated for 2 h in secondary antibody diluted in blocking solution (1:1000 Goat-α-rabbit AlexaFluor488; 1:1000 Goat-α-mouse AlexaFluor568) and washed again in PBS (4 × 10 min). Finally, sections air-dried at RT for 1 h before coverslipping with mounting medium containing [DAPI](https://www.sciencedirect.com/topics/neuroscience/dapi) (Prolong Gold; Thermofisher).

**Confocal image acquisition and analysis**

Confocal images of the CA1 region of the hippocampus for quantification of synaptic puncta were acquired at the Wohl Cellular Imaging Centre using an Inverted Spinning Disk confocal microscope (Nikon, Japan) and 60x oil immersion lens objective (NA 1.4). Exposure time was kept constant for the entire dataset. Images were 102.65 × 102.65 μm in size (512 × 512 pixels), acquired as a stack spanning 6−10 μm, at an interval of 0.3 μm. Three 3 non-continuous slices were imaged and analysed from each animal, and synaptic data averaged to a single datum for each animal. Synaptic puncta were analysed in ImageJ (<https://imagej.net/Welcome>), using a previously published pipeline (Half et al.). Analysis of synaptic puncta was performed in the strata oriens and radiatum – corresponding to the basal and apical dendritic regions respectively of CA1 pyramidal neurons **[Figure 2A]** and limited to 50x100µm Region of interest (ROI) 20µm either side of the stratum pyramidale to limit synaptic analysis to secondary and higher dendritic branching **[Figure 2A]**. In brief, 3 non-consecutive sections were manually selected based on quality of staining and contrast in the image. After background correction, images were thresholded. Thresholded images were used as a mask to measure puncta count: puncta smaller than 0.1 μm^2^ and larger than 4 μm^2^ were excluded top reduce noise in pouncta count (Half et al.). In all sections, GluA1 or bassoon expression was first determined. Synaptic GluA1 expression, defined as GluA1 puncta co-localized with bassoon, was determined by assessing of the number of GluA1 puncta that overlapped with bassoon puncta.

**Behaviour Data Analysis**

Videos were collected from all sample and test phases and analyzed for both social (sniffing stimuli, digging/burying near stimuli, etc.) and non-social (horizontal movement, vertical non-investigative behaviour, grooming, etc.) behaviours(3–9) using The Observer Video Analysis software (Noldus Information Technology, Wageningen, The Netherlands). Active sniffing within 1-2mm of a stimulus mouse-containing cylinder was considered social investigative behaviour. An investigation ratio was calculated; IR = N/(N+F), in which N is the time spent investigating the novel (in sample phases, N is the stimulus that will be replaced) and F is the time spent investigating the familiar stimulus. Sample phases typically have an investigation ratio of approximately 0.5, which is equivalent to chance. If experimental mice recognize a novel social stimulus, the investigation ratio is found to be statistically greater during the test than sample phase (3). Data points were removed when the total investigation time during test was lower than 5% of total test phase duration (<12 seconds in “easy” paradigm, <15 seconds in “difficult” paradigm), when only one stimulus was investigated during the sample or test phase, or when IR_Test_ values fell outside of 2 standard deviations from the mean (total n=19 across all experiments). Investigation ratios for sample phases were averaged for analysis.

**Statistical Analysis**

The arcsin transformed investigation ratios were analyzed with a mixed-design repeated measures ANOVA with treatment as the main factor and paradigm session (average sample and test) as the repeated-measure dependent variable. To reduce type I errors, specific mean comparisons were planned *a priori* to assess differences between IR_Sam_ and IR_Test_ within each treatment group. Specifically, paired t-tests were used to assess differences between IR_Sam_ and IR_Test_ within each treatment group. Additionally, one-way ANOVAs were used to assess treatment effects on IR_Test_, followed by Tukey *post hoc* tests. The durations of the other behaviours (Supplementary Table 1) were analyzed using a mixed-design repeated measures ANOVA with treatment as the main factor and the session of the paradigm (each of the sample phases and test) as the repeated-measures dependent variable, followed by Tukey *post hoc* tests. When normality failed, Kruskal-Wallis ANOVAs were performed followed by Dunn’s *post hoc* tests. One-way ANOVAs with Tukey *post hoc* tests were used to determine effects of treatment and task performance on GluA1 and bassoon puncta and GluA1/bassoon colocalization. SPSS and GraphPad Prism (v9.3.0) were used for all statistical analyses. Cohen’s d and eta-squared effect sizes are provided where appropriate. Two-tailed statistical significance was set at p<0.05.

1. Byers SL, Wiles MV, Dunn SL, Taft RA (2012): Mouse estrous cycle identification tool and images. *PloS One* 7: e35538.

2. Paxinos G, Franklin K (2001): *The Mouse Brain in Stereotaxic Coordinates*, 2nd ed. San Diego, CA: Academic Press.

3. Phan A, Lancaster KE, Armstrong JN, MacLusky NJ, Choleris E (2011): Rapid effects of estrogen receptor α and β selective agonists on learning and dendritic spines in female mice. *Endocrinology* 152: 1492–1502.

4. Phan A, Gabor CS, Favaro KJ, Kaschack S, Armstrong JN, MacLusky NJ, Choleris E (2012): Low doses of 17β-estradiol rapidly improve learning and increase hippocampal dendritic spines. *Neuropsychopharmacol Off Publ Am Coll Neuropsychopharmacol* 37: 2299–2309.

5. Phan A, Suschkov S, Molinaro L, Reynolds K, Lymer JM, Bailey CDC, *et al.* (2015): Rapid increases in immature synapses parallel estrogen-induced hippocampal learning enhancements. *Proc Natl Acad Sci U S A* 112: 16018–16023.

6. Sheppard PAS, Asling HA, Walczyk-Mooradally A, Armstrong SE, Elad VM, Lalonde J, Choleris E (2021): Protein synthesis and actin polymerization in the rapid effects of 17β-estradiol on short-term social memory and dendritic spine dynamics in female mice. *Psychoneuroendocrinology* 105232.
