## Supplementary Table for "Social memory in female mice is rapidly modulated by 17β-estradiol through ERK and Akt modulation of synapse formation"

**Supplementary Table 1**

List and description of mouse behaviours recorded during the social recognition paradigm. Based on (1).

| **Behaviour** | **Description** |
| --- | --- |
| **Stretch Approach** | Stretching towards stimulus with hind paws planted |
| **Sniff Stimulus** | Sniff/Investigation of stimulus, i.e. social investigation |
| **Bite Stimulus** | Biting cylinder or objects |
| **Dig** | Moving of bedding backwards with forepaws |
| **Bury** | Moving of bedding forwards with forepaws |
| **Horizontal Activity** | Includes walk, explore, sniff that does not fall into any of the above categories |
| **Vertical Activity** | Rear and lean on wall, lid sniff, lid chew (two paws on the floor of the cage), single jump |
| **Inactivity** | Sit, laydown, sleep, freeze |
| **Self-Groom** | Self-groom and scratch |
| **Stereotypies** | Strange behaviours: spinturns, repeated jumps, repeated lid chews (>3), head shakes, etc. |
| **Non-social Investigation** | Non-social sniffing of cylinder (above holes) |

1. Phan A, Lancaster KE, Armstrong JN, MacLusky NJ, Choleris E (2011): Rapid effects of estrogen receptor α and β selective agonists on learning and dendritic spines in female mice. *Endocrinology* 152: 1492–1502.
